# Missense variants in GABA_A_ receptor beta2 subunit disrupt receptor biogenesis and cause loss of function

**DOI:** 10.1101/2025.03.09.642292

**Authors:** Xi Chen, Ya-Juan Wang, Ting-Wei Mu

**Affiliations:** Department of Physiology and Biophysics, Case Western Reserve University School of Medicine, Cleveland, Ohio 44106, USA

**Keywords:** GABA_A_ receptor, GABRB2, epilepsy, missense variants, proteostasis, endoplasmic reticulum, folding, assembly, trafficking, degradation

## Abstract

Gamma-aminobutyric acid type A receptors (GABA_A_Rs) are the major inhibitory neurotransmitter-gated channel in the mammalian central nervous system. GABA_A_Rs function as heteropentamers, typically composed of two α1, two β2, and one γ2 subunits. Protein homeostasis between GABA_A_R folding, trafficking, assembly, and degradation is critical to ensure normal physiological functions. Variants in genes encoded for GABA_A_Rs lead to numerous neurological disorders, such as genetic epilepsy with or without neurodevelopmental delay. While these variants are associated with severe clinical presentations of epilepsy, the molecular mechanisms driving the disease remain to be elucidated. In this study, we focused on four missense epilepsy-associated variants (EAVs) in the *GABRB2* gene: Q209F210delinsH (c. 627_629del), R240T (c. 719G>C), I246T (c. 737T>C), and I299S (c. 896T>G). HEK293T cells exogenously expressing these β2 variants exhibited significantly reduced GABA-induced peak chloride current, indicating their loss of function. However, the four β2 EAVs differed in the degree of proteostasis deficiencies, including increased ER retention, compromised assembly, decreased protein stability, and reduced trafficking and surface expression, with Q209F210delinsH and R240T variants leading to the most severe degradation. Collectively, these results indicate that these epilepsy-linked variants have debilitating effects on the early biogenesis of the β2 subunit, causing misfolding, aggregation, and rapid degradation before it can be assembled with other subunits and transported to the plasma membrane. Overall, our work offers crucial mechanistic insight into how specific β2 missense variants impact the proteostasis maintenance of GABA_A_Rs, which could facilitate the development of effective therapeutics for genetic epilepsy by targeting trafficking-deficient GABA_A_R variants.

## Introduction

Gamma-aminobutyric acid type A receptors (GABA_A_Rs) are ligand-gated anion channels that mediate fast synaptic inhibition in the mammalian central nervous system. They play a critical role in maintaining the excitation-inhibition (E-I) balance and regulating neuronal activity (1). GABA_A_Rs belong to the Cys-loop family of receptors and are heteropentamers that assemble from a combination of 19 different subunits, including α(1–6), β(1–3), γ(1–3), δ, ε, π, θ, and ρ(1–3) (2). The most common type of synaptic receptors consists of two α1, two β2, and one γ2 subunits that are arranged in α-β-α-β-γ clockwise direction when viewed from the synaptic cleft (3, 4) (**Fig. 1A**). Each subunit contains shared structural elements: a large extracellular N-terminal domain (NTD), four transmembrane helices (TM1-4), a short intracellular TM1-2 loop, a short extracellular TM2-3 loop, a long intracellular TM3-4 loop, and a short extracellular C terminus. Two molecules of GABA bind at the interface between β and α subunits, which allosterically results in receptor activation in the transmembrane domain and allows the passage of the chloride ions through the channel.

**Figure 1.**
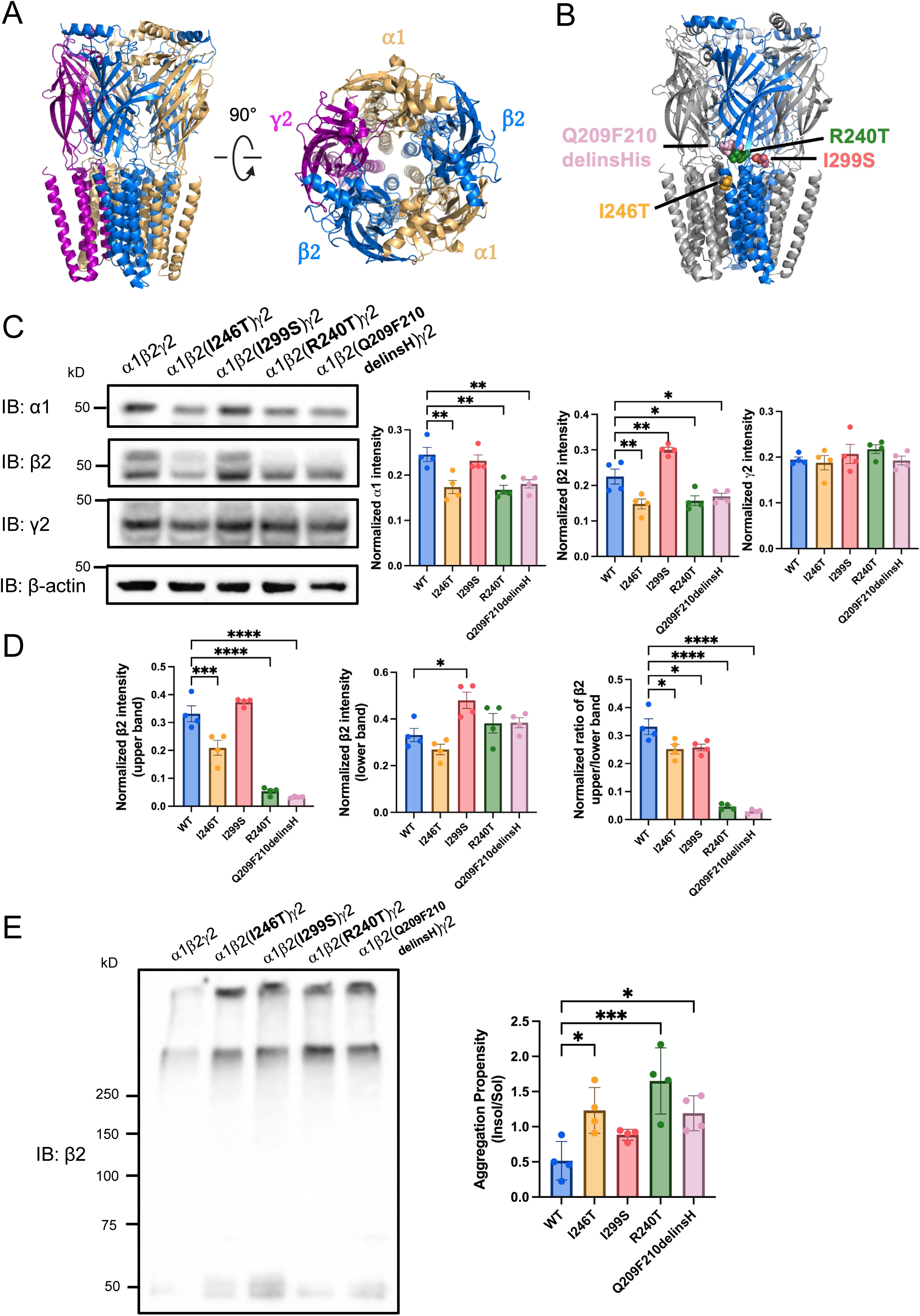
β2 missense epilepsy-associated variants (EAVs) in different region of the subunit lead to differing total protein expression and increased tendency to aggregate. (A) Cartoon representation of the pentameric GABA_A_ receptor (GABA_A_R), consisting of α-β-α-β-γ subunit assembled in clockwise direction when viewed from the synaptic cleft. PDB: 6X3S. (B) Structure of GABA_A_R, highlighting the positions of selected four missense variants in β2 subunit. Q209F210delinsH and R240T are located in the N-terminal domain near the entrance of the first transmembrane (TM) helix; I246T situates in the beginning of TM1; and I299S is in the extracellular TM2-3 loop region. (C) HEK293T cells transiently expressing β2 missense variants with WT α1 and WT γ2 (1:1:1; 0.25 µg each) were harvested forty-eight hours post transfection. Cells were lysed and solubilized in 2 mM n-dodecyl-β-maltoside (DDM). Total proteins were then subjected to SDS-PAGE and western blot to visualize the expression of α1, β2 and γ2 subunits. β2 protein displayed two bands at 55 and 48 kDa, respectively, which represented distinct glycoforms. β-actin was used as a loading control. Band intensity was quantified with ImageJ and normalized to the loading control, which is shown on the right (n=4). (D) Quantification of individual β2 glycoform band and the ratio of upper to lower band from (C). (E) Aggregation propensity of β2 EAVs. Cell pellet after extraction of soluble proteins (i.e., detergent-insoluble fraction) was washed and resuspended in Laemmli sample buffer containing 2% SDS and proceeded with western blot analysis. Band intensity of the entire lane was quantified, as large aggregates are seen at molecular weight above 250 kD. Insoluble-to-soluble-β2 ratio was calculated as a measure of aggregation propensity, which is shown on the right (n =4). Data is presented as mean ± SEM. One-way ANOVA followed by a post-hoc Dunnett’s test was used to determine statistical significance. * *p*< 0.05; **, *p* < 0.01; ***, *p* < 0.001; ****, *p* < 0.0001.

To become functional, GABA_A_R subunits must first fold into their proper three-dimensional structure with the aid of molecular chaperones and folding enzymes, then assemble with other subunits to form a pentamer in the endoplasmic reticulum (ER) (5). These correctly assembled heteropentamers further traffic through the Golgi apparatus to the plasma membrane (6–9). Misfolded or unassembled subunits will remain in the ER for additional folding/assembling cycles, and terminally misfolded proteins will be removed from the ER via cellular degradation pathways such as ER-associated degradation (ERAD) pathway in the proteasome (10, 11) or targeted to the lysosome via autophagy (12–15).

Given the essential role of GABA_A_Rs in inhibitory neurotransmission, variations in GABA_A_Rs that lead to receptor dysfunction upset the E-I balance, often resulting in neurodevelopmental and neuropsychiatric disorders such as autism, epilepsy, schizophrenia, and bipolar disorder (16–19). Currently, over 1000 clinical variants in genes encoding GABA_A_ receptor subunits have been reported in ClinVar (www.ncbi.nlm.nih.gov/clinvar/), including variants identified in key subunits such as α1 (20, 21), β2 (22–24), β3 (25, 26), and γ2 (27–31). Importantly, over 200 clinical variants have been identified in the β2 subunit, which contribute to various neurological disorders ranging from mild febrile seizures to severe epileptic encephalopathy (22–24). A recent study predicted the pathogenicity of these variants using ClinVar data and state-of-the-art computational tools, including AlphaMissense and Rhapsody (32), yet many reported variants have an uncertain significance due to a lack of experimental data. While previous studies characterized the phenotype and functionality of some of these epilepsy-associated β2 variants (22, 28), very little is known about how these variants affect the proteostasis regulation of the GABA_A_R, including folding, assembly, trafficking, and degradation. Proteostasis deficiency is one major disease-causing mechanism for GABA_A_ variants since many variants fail to reach the plasma membrane, rendering their resistance to current anti-seizure drugs on the market, which only act on the functional ion channels on the cell surface. Therefore, it is critical to characterize the proteostasis deficiency of GABA_A_ receptor variants, which paves the foundation for further mechanism-based therapeutic intervention.

In this study, we focus on four missense variations in the β2 subunit of GABA_A_R (encoded by the *GABRB2* gene): Q209F210delinsH (c. 627_629del), R240T (c. 719G>C), I246T (c. 737T>C), and I299S (c. 896T>G), which have been reported to be associated with severe epileptic phenotypes in patients. We demonstrated that all β2 variants caused a substantial reduction of GABA-induced peak chloride current, indicating a loss-of-function feature. Importantly, Q209F210delinsH, R240T, and I246T exhibited reduced steady-state protein expression, accelerated degradation kinetics, diminished ER-to-Golgi trafficking, and impaired receptor assembly, ultimately resulting in highly decreased surface protein expression. These results indicate that early proteostasis defects contribute to the dysfunction of the receptor. In contrast, the I299S variant showed similar protein stability, trafficking, assembly, and surface expression compared to the wild type (WT), suggesting channel gating defects rather than proteostasis dysregulation. Overall, our work fills in the knowledge gap about the proteostasis defects of these β2 variants, thereby enhancing our understanding of the underlying pathogenic mechanisms.

## Results

### Select ***β***2 missense variants lead to molecular defects and severe epileptic phenotypes

We selected four β2 missense variants that are located in various regions of β2 subunit and are associated with distinct phenotypical characteristics (**Fig. 1B**, **Table 1**). R240T and Q209F210delinsH are in the NTD, I246T variant is near the beginning of transmembrane helix 1 (TM1), and I299S is situated in the TM2-3 loop region that is known to be crucial for the channel gating process (32). From a biochemical perspective, replacing arginine with threonine at position 240 (R240T) eliminates the positive charge and disrupts the formation of the important salt bridge that helps stabilize the overall 3D structure. Q209F210delinsH, an indel variant that results in the deletion of two amino acids (Glu and Phe) and insertion of a new amino acid (His), will likely result in disturbed hydrophobic interactions from an added positive charge. Substituting isoleucine with threonine (I246T) introduces an additional hydroxyl group that can participate in hydrogen bonding, which can destabilize the hydrophobic protein core. The I299S variant may perturb the coupling between the agonist binding and the channel gating. Importantly, all of the missense variants were predicted to be “deleterious” with PolyPhen-2. suggesting that these amino acid substitutions have profound and harmful effects on the overall structure and/or function of the receptor, thus contributing to the epileptic phenotypes in patients carrying these variants.

**Table 1.**
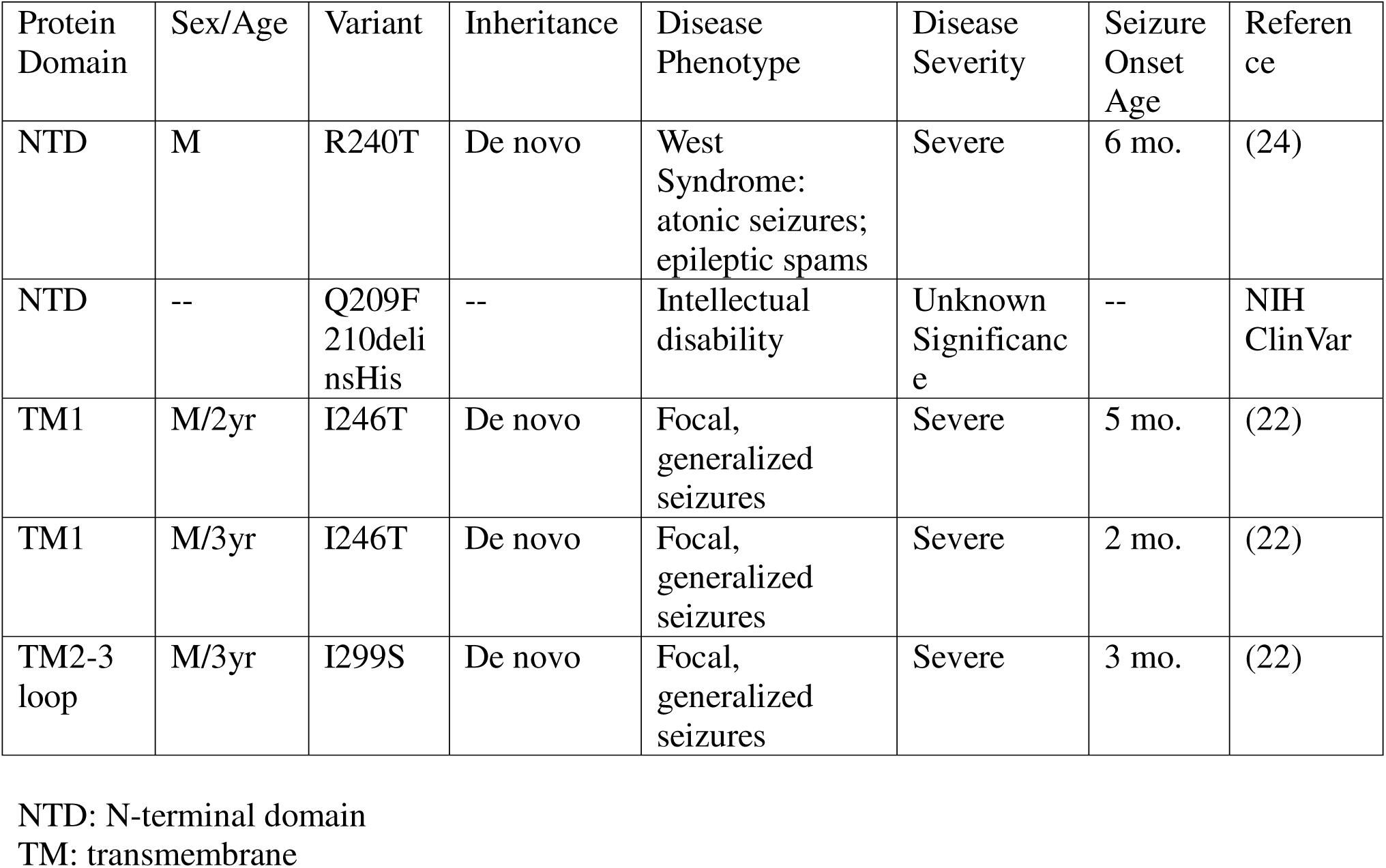
Clinical characteristics of GABA_A_R β2 missense variants (Q209F210delinsHis, R240T, I246T, and I299S).

| Protein Domain | Sex/Age | Variant | Inheritance | Disease Phenotype | Disease Severity | Seizure Onset Age | Reference |
| --- | --- | --- | --- | --- | --- | --- | --- |
| NTD | M | R240T | De novo | West Syndrome: atonic seizures; epileptic spasms | Severe | 6 mo. | (24) |
| NTD | -- | Q209F210delinsHis | -- | Intellectual disability | Unknown Significance | -- | NIH ClinVar |
| TM1 | M/2yr | I246T | De novo | Focal, generalized seizures | Severe | 5 mo. | (22) |
| TM1 | M/3yr | I246T | De novo | Focal, generalized seizures | Severe | 2 mo. | (22) |
| TM2-3 loop | M/3yr | I299S | De novo | Focal, generalized seizures | Severe | 3 mo. | (22) |

NTD: N-terminal domain
TM: transmembrane

**Table 1** summarizes the currently known clinical phenotypes of these missense variants. Two male individuals were reported to carry the I246T variant, one of which was deceased at age 2 years from respiratory failure in the setting of pneumonia (22). These patients had early seizures onset starting at age 2 and 5 months, respectively, and displayed epileptic syndromes consistent with early myoclonic encephalopathy (EME) or Lennox–Gastaut syndrome (LGS). Additionally, they were presented with severe developmental disorders, accompanied by either movement disorder or microcephaly. The male patient containing the β2(I299S) variant had early seizure onset age at 3 months. and demonstrated infantile spasms with severe developmental disorder, choreoathetosis, and microcephaly. R240T variant was identified in a male patient diagnosed with West syndrome, presenting with severe atonic seizures and epileptic spasms that began at 6 months of age, along with developmental delay (24). All of the variants mentioned above were de novo and linked with severe epileptic phenotypes, highlighting the need for further investigation into their disease-causing mechanisms on a molecular level. NIH ClinVar database recorded over 500 GABA_A_R β2 missense, frameshift, intron, or nonsense variants and classified them as benign, pathogenic, or uncertain significance. According to ClinVar, the Q209F210delinsH variant has been associated with intellectual disability, yet its precise pathogenicity is still unclear due to a lack of submissions. To date, there has been limited research on how these variants in the β2 subunit affect the biogenesis of the receptor. Therefore, investigating the underlying pathogenic mechanisms of these four β2 missense variants provides valuable insight into how the functional defect may contribute to the observed clinical phenotypes, potentially guiding the development of treatment strategies for patients afflicted by debilitating seizures.

### Select GABA_A_R ***β***2 missense variants demonstrate distinct steady-state protein levels and aggregation propensities

One of the major pathogenic mechanisms is that GABA_A_R epilepsy-associated variants (EAVs) are generally unable to fold into their native structure or assemble with other subunits, leading to trafficking deficiencies, excessive protein degradation, and impaired function (10, 33–35). Therefore, we first investigated how these missense variants in the β2 subunit affect the total and functional surface expression. β2 EAVs displayed variable total β2 protein levels in HEK293T cells co-transfected with equal amount of α1, β2 and γ2 subunit (**Fig. 1C**). I246T, R240T, and Q209F210delinsH variants showed around 30% reduction in β2 protein expression with equal β2 mRNA levels (**Fig. 1C, Supplemental Fig. S1**), suggesting the post-transcriptional effect of such variants. Interestingly, the I299S variant led to slightly increased β2 protein levels compared to the WT (**Fig. 1C**). Because we co-transfected all three subunits to form functional heteropentamers, we also examined the total protein expression levels of α1 and γ2 in these β2 EAVs. The α1 protein levels in HEK293T cells carrying I246T, R240T, and Q209F210delinsH β2 variants were significantly decreased despite the same amount of α1 being transfected (**Fig. 1C)**. The diminished α1 protein levels in these β2 variants suggest that the presence of β2 missense variants has a negative effect on the proteostasis of α1 subunit, possibly due to the increased degradation of the unassembled α1 subunit. Interestingly, total γ2 protein levels were less affected by the presence of β2 variants (**Fig. 1C**).

Notably, β2 subunit, which contains three N-linked glycosylation sites at Asn32, Asn104, and Asn173, has a migration pattern on a gel as two distinct bands at 48 kDa and 55 kDa, representing different β2 glycoforms (**Fig. 1C**). While the lower band intensity of I246T, R240T and Q209F210delinsH variants showed no difference, the intensity of the upper band, which is the mature and fully glycosylated β2 protein, was significantly reduced in these variants (**Fig. 1C**, **1D**), suggesting their potential glycosylation defect. Notably, intensities of both upper and lower band of I299S variant were increased compared to WT. Overall, all the selected β2 EAVs demonstrated a reduced ratio of upper band intensity to lower band intensity (**Fig. 1D**).

Additionally, we assessed the extent of protein aggregation using a detergent solubility assay. After lysing the cells and solubilizing cell lysates in n-dodecyl-β-maltoside (DDM), cell pellet was re-suspended in a buffer containing 2% SDS and 10% β-mercaptoethanol (β-ME) and visualized via SDS-PAGE and Western blot analysis. All β2 EAVs exhibited elevated β2 levels in the insoluble fraction compared to the WT, especially showing large aggregates at molecular weights exceeding 250 kDa (**Fig. 1E**). Aggregation propensity was further calculated as the ratio of insoluble (pellet) over soluble (supernatant) fraction. We observed that I246T, R240T, and Q209F210delinsH variants demonstrated a greater aggregation propensity than WT, with R240T showing the most significant change around 3.2-fold increase (**Fig. 1E**).

### Select GABA_A_R ***β***2 missense variants demonstrate distinct surface protein expression and highly diminished receptor function

Since GABA_A_Rs need to be transported to the cell surface to become functional, we examined the surface expression levels of select β2 EAVs using a surface biotinylation assay. Surface proteins were labeled with membrane-impermeable Sulfo-NHS SS-Biotin on ice and affinity-purified with neutravidin-conjugated beads, which were then analyzed using SDS-PAGE and Western blot technique. Surface α1, β2 and γ2 protein levels in I246T variant were significantly reduced by ∼45%, indicating trafficking defect and impaired receptor assembly process (**Fig. 2A**). The two NTD variants, R240T and Q209F210delinsH, demonstrated the most dramatic decrease in surface expression of α1, β2 and γ2, with a decline of over 90%. However, I299S variant seemed to have little impact on the surface expression of α1 and γ2 with a slight but significant increase in surface β2 level (**Fig. 2A**). Using non-permeabilizing immunocytochemistry staining, we observed a similar decline in surface β2 protein intensities for I246T, R240T and Q209F210delinsH EAVs, with no change in surface β2 intensity for I299S variant (**Fig. 2B**). Overall, surface protein expression data clearly demonstrated that β2 I246T, R240T and Q209F210delinsH variants led to significantly reduced surface trafficking of α1, β2, and γ2 subunits, possibly due to their early protein biogenesis defects, including the misfolding of β2 variants and subsequent subunit assembly deficiencies.

**Figure 2.**
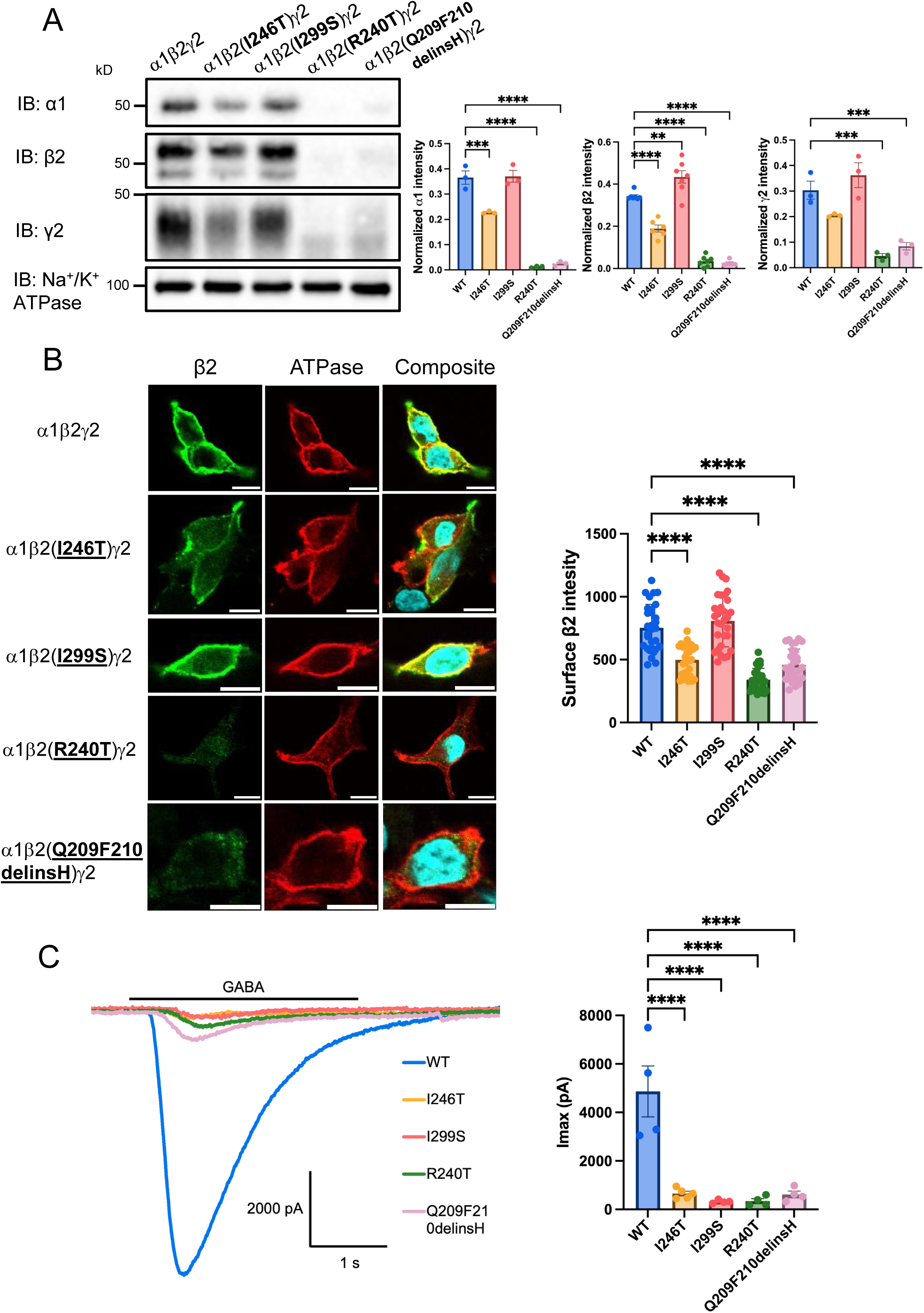
Epilepsy-associated missense variants in. β**2 subunit impair receptor functional surface expression.** (**A**) Surface expression of β2 EAVs using cell surface biotinylation assay. HEK293T cells transiently expressing β2 EAVs were incubated with Sulfo-NHS SS-biotin on ice to label membrane proteins and lysed forty-eight hours post transfection. Cell lysates containing biotinylated proteins were affinity-purified with neutravidin-conjugated beads and subjected to SDS-PAGE and western blot to visualize the expression of surface α1, β2 and γ2 proteins. Na^+^/K^+^ ATPase was used as a loading control. Quantification of band intensity normalized to the loading control is shown on the right (n=3 or 7). (**B**) Surface β2 intensity using immunofluorescence staining. HEK293T cells expressing β2 EAVs were fixed with 4% paraformaldehyde without membrane permeabilization and incubated with anti-GABA_A_R β2/3 antibody and anti-Na^+^/K^+^ ATPase antibody. The fluorescence intensity of surface β2 was assessed with ImageJ and the quantification is shown on the right (scale bar = 20 µm, n=30). (**C**) GABA-induced peak chloride current of β2 EAVs using automated patch clamping electrophysiology. GABA was applied at a saturating concentration of 100 µM for 3 seconds as shown by the horizontal bar above the current traces. Whole-cell currents were recorded using the IonFlux Mercury 16 ensemble plates at a holding voltage of −60 mV. Representative whole-cell voltage-clamp recording traces are shown. Quantification of the peak currents (I_max_) is shown on the right (n =4). pA: pico Ampere. Data is presented as mean ± SEM. One-way ANOVA followed by a post-hoc Dunnett’s test was used to determine statistical significance. **, *p* < 0.01; ***, *p* < 0.001; ****, *p* < 0.0001.

Given the close relationship between surface expression and function, we further investigated GABA-induced maximum chloride current using automated patch-clamping electrophysiology. HEK293T cells transiently transfected with WT or variant β2 together with α1 and γ2 subunits were exposed to a saturating concentration of GABA (100 µM) for 3 seconds, and the current amplitudes were recorded using whole-cell patch-clamping technique. All selected β2 EAVs showed highly decreased peak current amplitudes upon GABA application (**Fig. 2C**), indicating a loss-of-function phenotype. Compared to WT receptor that displayed an average peak current amplitude of 4.87 ± 1.05 nA, I246T, I299S, R240T, and Q209F210delinsH EAVs exhibited average peak current amplitudes of 0.66 ± 0.09 nA, 0.32 ± 0.03 nA, 0.34 ± 0.10 nA, and 0.62 ± 0.14 nA, respectively. Our data is consistent with previous electrophysiological studies, which showed that I246T variant had a substantially reduced GABA-induced current compared to WT (22).

Overall, I246T, R240T, and Q209F210delinsH EAVs demonstrated reduced total and surface expression, indicating impaired proteostasis regulation that involves trafficking and assembly. Reduced surface expression of these variants further correlated with the diminished receptor function. Interestingly, while I299S variant had comparable total and surface expression levels as the WT, its ability to conduct current was highly compromised. Since the TM2-3 loop is critical for channel gating (32), the loss-of-function trait observed in I299S variant can be attributed to its gating defects, rather than issues in the early biogenesis pathway.

### GABA_A_R ***β***2 EAVs demonstrate differing protein stability and degradation preferences

Since select GABA_A_R β2 EAVs exhibited a decrease in their total and surface protein expression levels, these variants could be rapidly removed via either proteasomal or lysosomal degradation pathways. First, we determined the protein stability and steady-state protein levels of the β2 variants using a cycloheximide-chase assay. HEK293T cells transiently expressing WT receptor or β2 EAVs were treated with cycloheximide to inhibit protein synthesis. Compared to the WT, the two NTD variants R240T and Q209F210delinsH demonstrated a significant reduction in protein levels at 2, 3, and 4 hours post application of cycloheximide, indicating a decreased protein stability (**Fig. 3A**). Similarly, protein expression of I246T is significantly diminished at 6 hours after cycloheximide treatment compared to WT (**Fig. 3B**). Conversely, a six-hour cycloheximide assay showed no difference in degradation kinetics between WT and I299S variant (**Fig. 3B**). The varying protein stability and turnover rate corresponded to the total protein expression of β2 variants (**Fig. 1C**).

**Figure 3.**
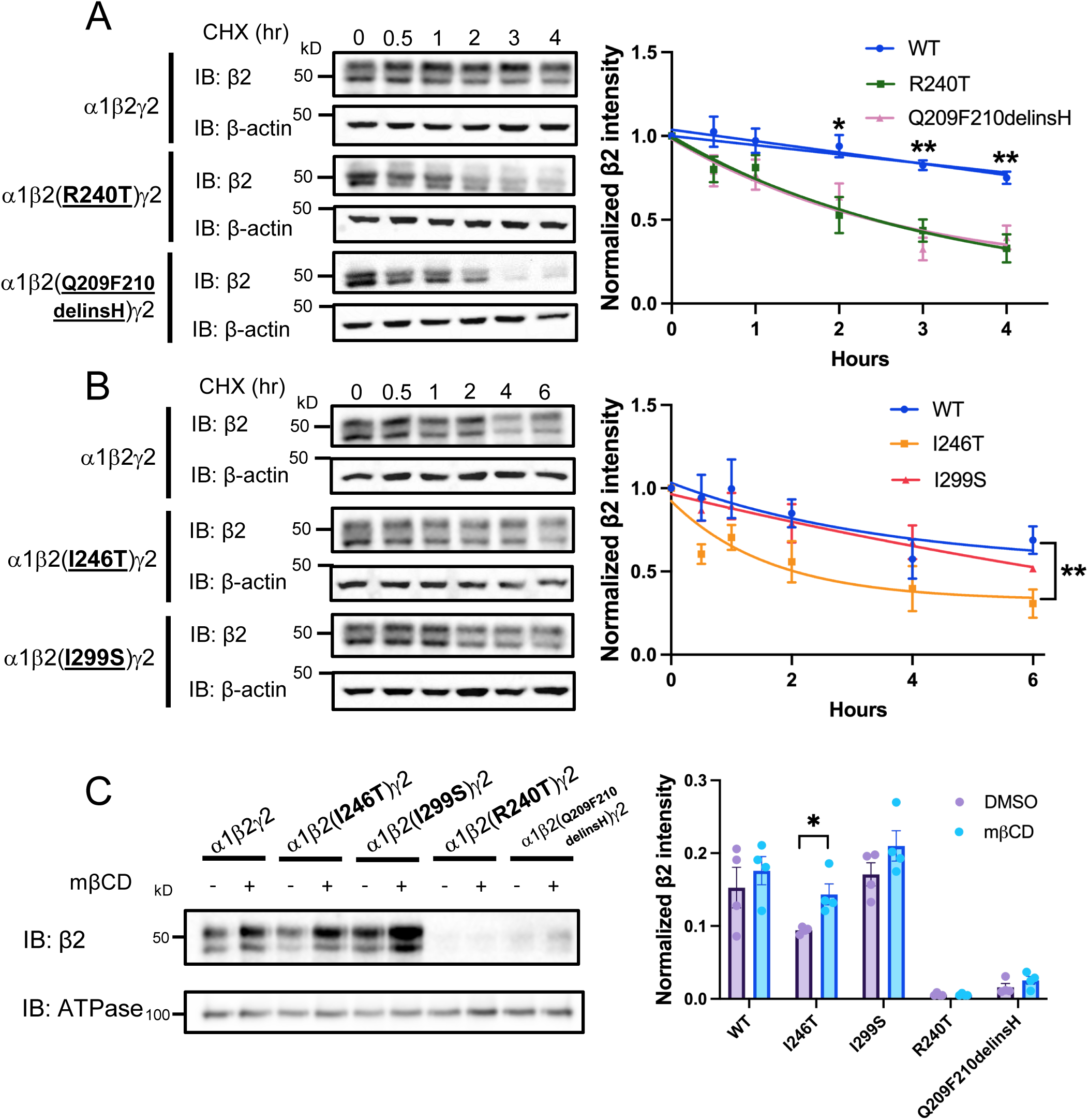
β**2 EAVs demonstrate distinct protein stability and varying levels of endocytosis.** To determine protein stability of β2, HEK293T cells expressing β2 EAVs were treated with cycloheximide (CHX, 100 µg/mL) for 0 to 6 hours to block protein synthesis. Total cell lysates were subjected to western blot to visualize the degradation kinetics of β2 protein. (**A**) A four-hour CHX assay was performed for WT, R240T and Q209F210delinsH (n=3 or 4). (**B**) A six-hour CHX assay was performed for WT, I246T, and I299S (n=3). A one-phase decay non-linear regression was used to fit the data using prism. Data is presented as mean ± SEM. One-way ANOVA followed by a post-hoc Dunnett’s test was used to determine statistical significance between WT, I246T and I299S, and between WT, R240T, and Q209F210delinsH at individual time point. (**C**) To determine the extent of endocytosis for β2 variant receptors, HEK293T cells expressing β2 EAVs were treated with methyl-β-cyclodextrin (mβCD, 100 µM), which removes cholesterol from the plasma membrane to inhibit clathrin-mediated receptor endocytosis. Surface β2 expression was determined with a cell surface biotinylation assay, followed by western blot analysis (n=4). A Student’s *t* test was performed to determine the statistical significance between DMSO and mβCD-treated condition for all β2 variants. * *p*< 0.05; **, *p* < 0.01.

Additionally, protein variants often exhibit reduced stability at the cell surface, which can trigger accelerated endocytosis. As a result, these variants are more likely to be internalized and subsequently degraded through the endosomal-lysosomal pathway (36). To assess protein stability on the surface, HEK293T cells expressing WT and β2 EAVs were treated with methyl-β-cyclodextrin (mβCD), which depletes cholesterol from the plasma membrane and inhibits clathrin-mediated endocytosis (37). Compared to WT that showed a 1.2-fold increase in surface expression upon mβCD treatment, I246T demonstrated a larger 1.5-fold increase, indicating a higher rate of endocytosis (**Fig. 3C**). I299S exhibited a similar endocytosis rate compared to WT, while R240T and Q209F210delinsH did not undergo substantial internalization, suggesting that their lack of surface expression is likely due to a trafficking defect rather than excessive endocytosis. Overall, I246T, R240T, and Q209F210delinsH variants showed reduced steady-state protein levels and elevated degradation rate, while I299S variant had similar stability and degradation kinetics as the WT. In addition, I246T displayed an enhanced endocytosis.

Misfolded proteins or unassembled subunits will be degraded by the ER-associated degradation (ERAD) pathway via the ubiquitin-proteasome machinery (38–40). Additionally, large aggregates can be cleared in the lysosome through autophagy (12, 41). To determine the preferential degradation pathways of β2 EAVs, HEK293T cells transiently expressing WT or β2 variants were treated with MG132 or bafilomycin (bafA1) for 6 hours to inhibit the proteasome or lysosome, respectively. Application of MG132 led to an increase in ubiquitinated protein levels, whereas bafA1 treatment resulted in an increased LC3bII/LC3bI ratio, indicating impaired proteasomal activity and autophagic flux, respectively (**Fig. 4A**). While WT β2 displayed a comparable increase when treated with either MG132 or bafA1, R240T variant exhibited a significantly greater increase in protein expression with MG132 treatment, compared to bafA1 treatment (**Fig. 4A**, **Fig. 4B**). Similar to WT, I246T and Q209F210delinsH variants appeared to utilize both lysosomal– and proteasomal-dependent degradation pathways efficiently. I299S variant, which demonstrated elevated total and surface protein expression, showed only a minimal increase in β2 expression following the addition of either MG132 or bafA1.

**Figure 4.**
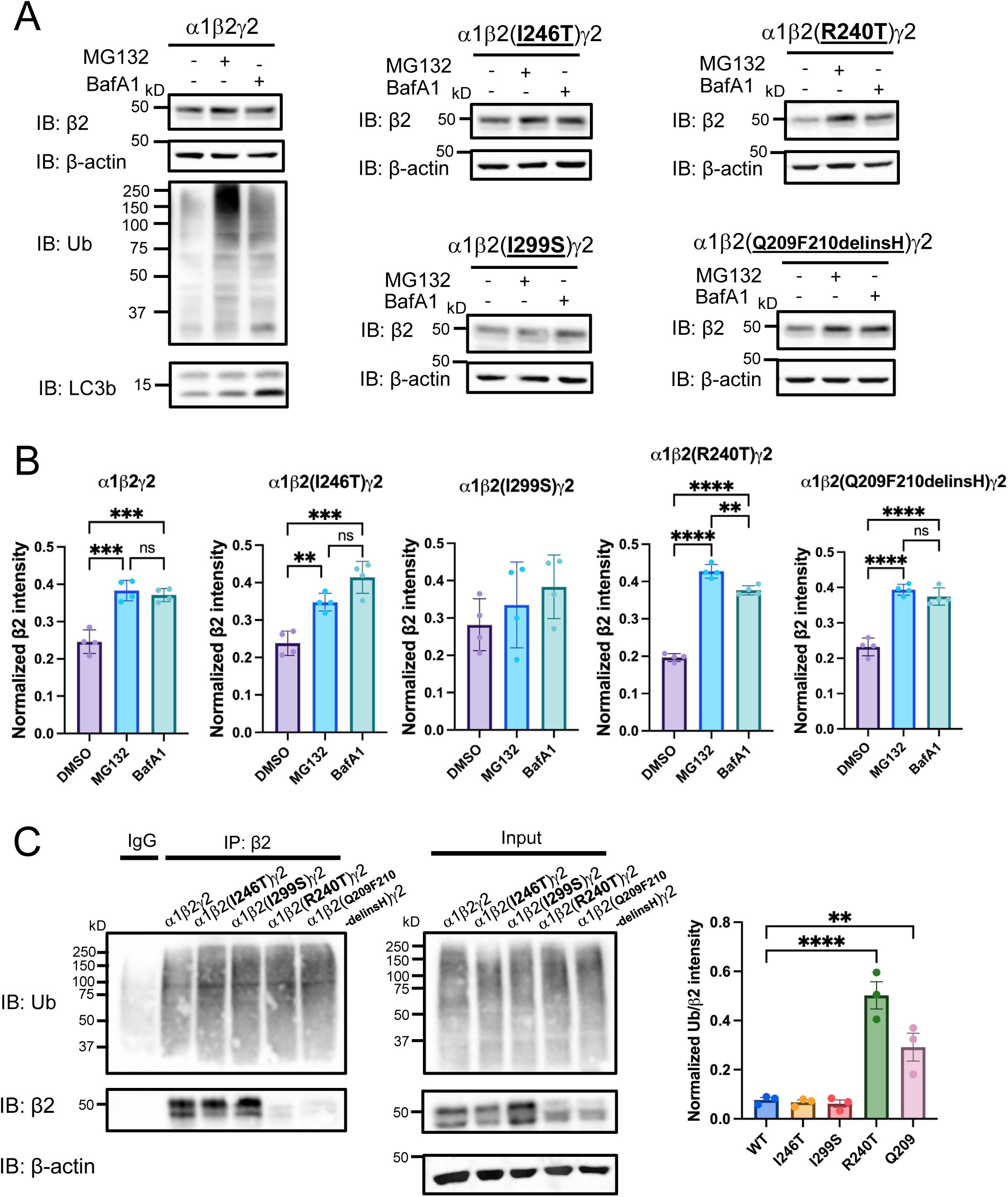
β**2 EAVs engage both proteasomal and lysosomal degradation pathways.** (**A**) To evaluate the degree of proteasomal and lysosomal degradation, HEK293T cells transiently expressing β2 EAVs were treated with MG132 (10 µM, 6 hrs) or bafilomycin A1 (bafA1, 1 µM, 6 hrs) to inhibit the proteasome or lysosome, respectively. Cells were lysed forty-eight after transfection and total protein was visualized using SDS-PAGE and western blot analysis (n=4). (**B**) Quantification of band intensity normalized to the loading control (β-actin). (**C**) Ubiquitination levels of β2 EAVs using a co-immunoprecipitation (co-IP) assay. HEK293T cells were co-transfected with HA-tagged ubiquitin along with WT or β2 EAVs. Cells were treated with MG132 (10 µM, 6 hrs) to inhibit proteasomal degradation and harvested forty-eight hours post transfection. 1000 µg of total lysates were pre-cleared and incubated with anti-GABA_A_R β2 antibody, followed by incubation with protein A/G beads to pull down β2 and β2-interacting proteins. Eluent from the beads were proceeded with SDS-PAGE and western blot to detect ubiquitin levels. Quantification of Ubiquitin, which is normalized to the immunoprecipitated β2, post co-IP is shown on the right (n=3). IP: immunoprecipitation. Data is presented as mean ± SEM. One-way ANOVA followed by a post-hoc Dunnett’s test was used to determine statistical significance. **, *p* < 0.01; ***, *p* < 0.001; ****, *p* < 0.0001; ns, not significant.

Furthermore, we performed a co-immunoprecipitation assay to examine the ubiquitination of β2. HEK293T cells co-expressing WT or β2 EAVs and HA-tagged ubiquitin were treated with MG132 to inhibit proteasomal degradation. Cell lysates were then co-immunoprecipitated with β2 antibody and probed for ubiquitin. The ratio of ubiquitin intensity to β2 post co-immunoprecipitation was quantified as a measure of β2 ubiquitination. We found that the two NTD rapidly degrading and less stable variants, R240T and Q209F210delinsH, displayed a significantly higher level of ubiquitinated β2 compared to WT (**Fig. 4C**). The increased ubiquitination of β2 is consistent with the results shown in **Fig. 4A** and **Fig. 4B**, indicating that R240T and Q209F210delinsH are heavily ubiquitinated and subjected to efficient cellular degradation.

### GABA_A_R ***β***2 EAVs show trafficking and assembly defects and are retained in the ER

Given the diminished surface expression levels of I246T, R240T, and Q209F210delinsH variants, these β2 variants are likely unable to assemble with other subunits and further anterograde traffic to the surface. To determine the trafficking efficiency of β2 variants, we performed an endoglycosidase H (endo H) digestion assay. Endo H cleaves simple glycans that are added in the ER, but is unable to cleave more complex glycans that are further modified in the Golgi. Peptide-N-Glycosidase F (PNGase F) cleaves almost all kinds of glycan and was used as a positive control for unglycosylated β2 (**Fig. 5A**, lane 3). We labeled bands that have the same molecular weight as unglycosylated β2 as endo H-sensitive bands, whereas bands at higher molecular weights were labeled as endo H-resistant. The ratio of endo H-resistant to total β2 bands was quantified as a measure of ER-to-Golgi trafficking efficiency. β2 variants that demonstrated a decrease in surface expression (i.e., I246T, R240T, and Q209F210delinsH) also displayed a reduced trafficking efficiency (**Fig. 5A**, cf. lanes 5, 9, 11 to lane 2). Importantly, R240T and Q209F210delinsH showed 94% and 99% reduction in trafficking efficiency, which is consistent with their substantial decrease in surface expression (**Fig. 2A**). In contrast, I299S showed a similar ER-to-Golgi trafficking efficiency compared to the WT (**Fig. 5A**, cf. lane 7 to lane 2), which aligns with its surface expression (**Fig. 2A**).

**Figure 5.**
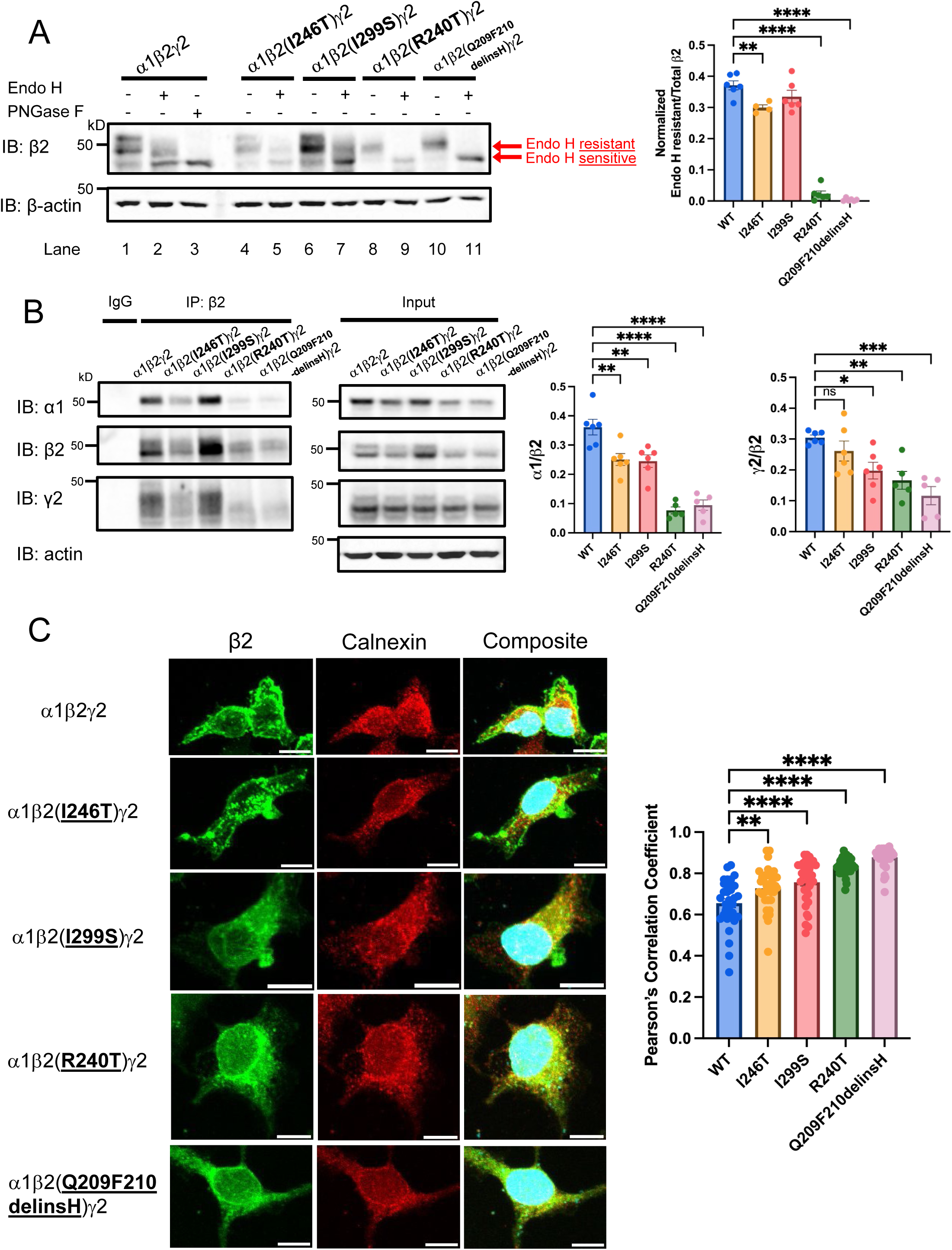
β**2 EAVs are ER-retained and exhibit deficiencies in trafficking and subunit assembly.** (**A**) ER-to-Golgi trafficking efficiency was determined using an endoglycosidase H (endo-H) digestion assay. Cleavage by Peptide-N-Glycosidase F (PNGase F) enzyme served as a control for unglycosylated β2 subunits (lane 3). After endo H digestion, β2 subunits with a molecular weight equal to PNGase F-treated band were endo H-sensitive, whereas those with higher molecular weights were endo H-resistant. The ratio of endo H-resistant β2 to total β2 bands was quantified to assess ER-to-Golgi trafficking efficiency, which is shown on the right (n=6). (**B**) Interaction between β2 and α1 and between β2 and γ2 was evaluated using a co-IP experiment. Quantification of the ratio of proteins of interest (i.e., α1 or γ2) to β2 post co-IP is shown in the right panels (n=6). IP: immunoprecipitation. (**C**) Immunofluorescence staining showing the colocalization of β2 (green) with calnexin (red), an ER resident chaperone. Pearson’s coefficient was obtained using ImageJ colocalization analysis and quantification is shown on the right (n= 32-34). Data is presented as mean ± SEM. One-way ANOVA followed by a post-hoc Dunnett’s test was used to determine statistical significance. * *p*< 0.05; **, *p* < 0.01; ***, *p* < 0.001; ****, *p* < 0.0001; ns, not significant.

Since individual subunits must assemble with other subunits to be properly transported to the cell surface, we next probed the subunit assembly process with a co-immunoprecipitation assay. Compared to the WT receptor, I246T, R240T and Q209F210delinsH variants demonstrated a decreased level of α1 subunit that co-immunoprecipitated with β2 subunit (**Fig. 5B, Supplemental Fig. S2A**). I246T and I299S variants exhibited a 30% reduction in α1-β2 interaction, as quantified by the ratio of α1 to β2 post co-immunoprecipitation, while R240T and Q209F210delinsH variants demonstrated a more drastic decline with average decreases of 80% and 75%, respectively. Similarly, there was a decreased interaction between β2 and γ2 subunit for all β2 variants, with R240T and Q209F210delinsH variants showing the most significant reduction (**Fig. 5B**, **Supplemental Fig. S2B**). These data suggests that β2 EAVs, particularly I246T, R240T, and Q209F210delinsH, exhibit impaired receptor assembly and insufficient trafficking, which contribute to a decreased functional surface expression.

Given the diminished ER-to-Golgi trafficking efficiency of β2 EAVs, we further confirmed the ER-retention of these variants using immunofluorescence staining. All selected β2 EAVs demonstrated an increased colocalization of β2 with calnexin, an ER-resident type I transmembrane protein, as quantified by Pearson’s Correlation Coefficient (**Fig. 5C**). Collectively, these results indicate that these epilepsy-associated variants (i.e., I246T, R240T, Q209F210delinsH) result in compromised subunit assembly, impaired receptor trafficking, reduced protein stability, and efficient removal via lysosomal or proteasomal degradation pathways. The overall disruption in proteostasis regulation ultimately contributes to diminished surface expression and a loss-of-function phenotype.

## Discussion

Molecular characteristics of the four β2 EAVs are summarized in **Figure 6**. All four β2 epilepsy-associated variants lead to highly reduced peak chloride current upon GABA application compared to the WT receptor (**Fig. 2C**), indicating their loss of function. However, their dysfunctions have different underlying causes. I299S variant, located in the TM2-3 loop region, showed similar protein stability, oligomerization, trafficking efficiency, and surface expression to the WT. Therefore, its functional impairment is likely not due to issues in the early biogenesis pathway, but rather due to deficiency in channel gating. As previous studies highlighted the critical role of the extracellular TM2-3 loop in coupling the agonist binding to receptor activation (32), future experiments are needed to more specifically investigate the disease-causing mechanisms of I299S, such as whether the variant alters the channel kinetics.

**Figure 6.**
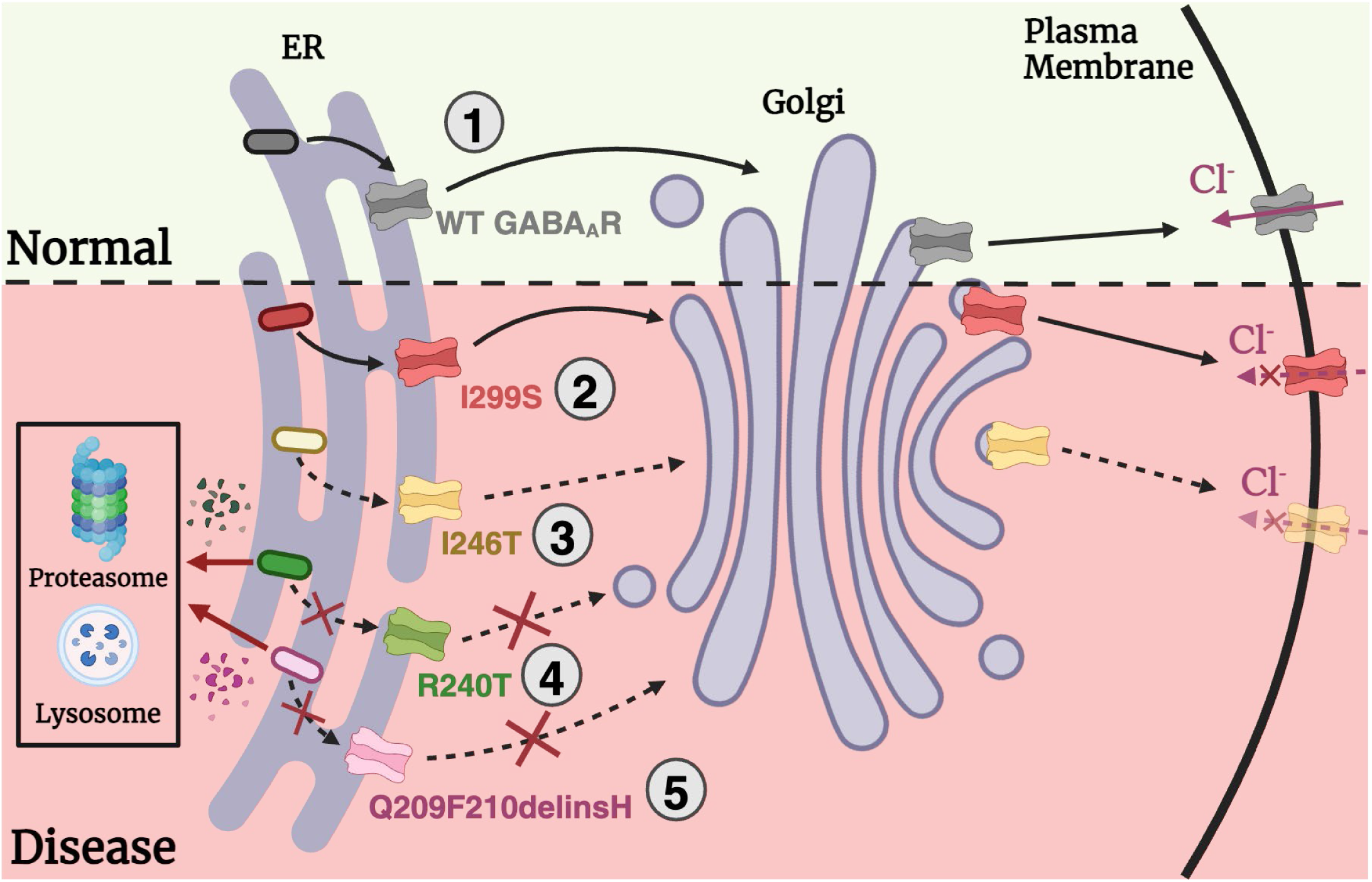
Overview of the proposed pathogenic mechanisms associated with GABA_A_R. β**2 EAVs**. Under normal physiological conditions, WT GABA_A_R β2 subunit folds and assembles with α1 and γ2 subunit. Fully assembled heteropentameric receptor is then trafficked to the plasma membrane, where it becomes a functional, anion-conducting channel (Case 1). However, in the disease state, β2 variants fail to create a functional receptor on the surface due to various underlying causes (Cases 2-5). I299S variant is able to reach the surface, but could not generate any ion flow, likely due to gating defects that impact channel kinetics (Case 2). I246T variant shows reduced efficiency in assembling and trafficking, reaching plasma membrane only about 50% of the time. Once on the surface, I246T is still unable to respond to high extracellular GABA, suggesting the presence of additional channel gating defects (Case 3). NTD R240T and Q209F210delinsH variants undergo substantial degradation via the proteasome or lysosome and are virtually incapable of assembling or trafficking. Consequently, they display a severe reduction in surface expression and could not conduct any current (Cases 4 and 5).

In contrast to I299S, TM1 variant (I246T) displayed reduced surface trafficking, insufficient subunit assembly, increased ER retention, and reduced protein stability. While its surface expression was decreased by 50%, GABA-induced peak chloride was reduced by 87% compared to WT. The 50% decrease in surface expression alone cannot account for the much more severe functional impairment, suggesting additional alterations in channel gating may be involved. Our result is consistent with previous electrophysiological studies, which found that I246T variant had reduced GABA-induced current compared to WT and a heightened agonist sensitivity at low GABA concentrations (22).

The two NTD variants, R240T and Q209F210delinsH, led to more severe phenotypes. These variants exhibited a significantly reduced ER-to-Golgi trafficking efficiency and impaired receptor assembly process. Additionally, they were retained in the ER, showed reduced protein stability, and efficiently utilized both lysosomal and proteasomal degradation pathways, ultimately leading to a near complete loss of functional surface expression. Their lack of surface expression corroborates with the substantial decrease in GABA-induced peak chloride current. Therefore, these two NTD variants (R240T and Q209F210delinsH) may be highly destabilizing, leading to protein misfolding and rapid degradation during the early biogenesis process. Interestingly, R240T variants also exhibited slight preference for proteasomal degradation and increased ubiquitination levels. A previous study found that GABA_A_R misfolding-prone α1(A322D) variant utilized ERAD pathway, involving Grp94 chaperone-mediated recognition, Hrd1-mediated ubiquitination, and VCP (valosin-containing protein)-mediated extraction (10). Destabilized R240T variant could engage similar degradation machinery, which warrants further investigation. HEK293T cells used in this study offer a unique advantage to study mechanisms of receptor biogenesis, including folding, trafficking, assembly, and degradation, as they do not express endogenous GABA_A_Rs and thus precise control of the subunit combination can be achieved. In addition, future experiments using induced pluripotent stem cell (iPSC)-derived cortical neurons (33, 42) or mice (43, 44) harboring these variants are important to validate these findings and gain deeper insight into the pathogenic mechanisms in neuronal environment.

Importantly, all selected β2 variants unanimously demonstrated increased aggregation propensity and ER retention. Protein aggregates can disrupt the normal cellular function and homeostasis. Excessive formation of aggregates, along with retention of these variants in the ER, can cause ER stress. Accumulation of misfolded protein can activate stress-adaptive unfolded protein response (UPR) pathway in the ER. UPR attempts to increase protein folding capacity and re-establish homeostasis through the action of three arms: 1) the ATF6 (activating transcription factor 6), 2) IRE1 (inositol-requiring enzyme 1), and 3) PERK (protein kinase R-like ER kinase) pathways (45–47). By causing ER stress, these β2 EAVs may activate different components of the UPR. Previous work found that GABA_A_R α1 frameshift variants activated the UPR to various degrees, leading to an upregulation of BiP or CHOP, which are signatures of ATF6 or PERK pathway activation (48).

Furthermore, UPR modulation can serve as an effective tool to mitigate proteostasis defects. Pharmacological activation of the UPR has been shown to increase the folding, trafficking, and functional surface expression of various misfolding-prone and trafficking-deficient GABA_A_R variants, including α1(A322D) and α1(D219N) variants (49, 50). Proteostasis regulators adapt the proteostasis network to improve ER well-being (51, 52). Examples of proteostasis regulators include FDA-approved drugs (i.e., suberoylanilide hydroxamic acid (SAHA), dinoprost, and dihydroergocristine), L-type channel blockers (i.e., verapamil), and BIX (a potent BiP activator), which improved the trafficking and function of variant-containing GABA_A_ receptors and NMDA receptors (34, 35, 53, 54). In addition, GABA_A_R-specific pharmacological chaperones such as hispidulin and TP003 were shown to enhance the membrane-activity of pathogenic GABA_A_R variants (33), and the combination of pharmacological chaperones and proteostasis regulators may yield further synergistic effects (55). While numerous anti-seizure drugs have been developed, ∼30% of patients suffering from genetic epilepsy are resistant to current drug treatment, partially due to lack of GABA_A_R surface expression (56). Therefore, there is a critical need to develop effective therapeutics to target defective GABA_A_ receptor variants to treat genetic epilepsy. Understanding cellular stress-adaptation mechanisms can offer valuable insight into therapeutic strategies for UPR activation to alleviate cellular stress, restore proteostasis, and ultimately rescue the function of these disease-associated GABA_A_R variants.

## Materials and methods

### Chemicals

GABA (catalog #: A2129) was purchased from Millipore Sigma. MG132 (ApexBio, catalog #: A2585) was used to inhibit proteasome at 10 µM for 6 hours. Bafilomycin A1 (BafA1; Cayman Chemical, catalog #: 11038) was used to inhibit lysosome at 1 µM for 6 hours. Cycloheximide (CHX; Enzo Life Sciences, catalog #: ALX-380-269) was used to inhibit protein synthesis at 100 µg/mL from 0-6 hours. methyl-β-cyclodextrin (mβCD; Ambeed, catalog #: A539125) was used to block receptor endocytosis at 100 µM for 18 hours.

### Plasmids

The pCMV6 plasmids from Origene include human GABA_A_ receptor α1 subunit (*GABRA1*) (Uniprot #: P14867-1) (catalog #: RC205390), β2 subunit (*GABRB2*) (isoform 2, Uniprot #: P47870-1) (catalog #: RC216424), γ2 subunit (*GABRG2*) (isoform 2, Uniprot #: P18507-2) (catalog #: RC209260). The missense variants in the GABA_A_ receptor β2 subunit, Q209F210delinsH, R240T, I246T, and I299S, were generated using QuikChange II site-directed mutagenesis kit (Agilent Genomics, catalog #: 200523), and the cDNA sequences were confirmed via DNA sequencing.

### Cell culture and transfection

HEK293T cells were obtained from Abgent (catalog #: CL1032) or ATCC (catalog #: CRL-3216, donor sex: female) and maintained in Dulbecco’s Modified Eagle Medium (DMEM) (Fisher Scientific, Waltham, MA, catalog #: SH3024301) with 10% heat-inactivated fetal bovine serum (Fisher Scientific, catalog #: SH30396.03HI) and 1% Penicillin-Streptomycin (Fisher Scientific, catalog #: SV30010) at 37 °C in 5% CO2. Monolayers were passaged upon reaching 90-95% confluency using 0.05% trypsin protease (Fisher Scientific, catalog #: SH30236.01). Cells were grown in 6-well plates, 6-cm, or 10 cm dishes and allowed to reach ∼50% confluency before transient transfection according to the manufacturer’s instruction. TransIT-2020 (Mirus Bio, Madison, WI, catalog #: MIR 5406) and Opti-MEM (ThermoFisher, catalog #: 31985070) were used for transient transfection according to the manufacturer’s instruction. Cells were harvested for experiments forty-eight hours post transfection.

### Western blot analysis

HEK293T cells were washed with DPBS (Fisher Scientific, catalog #: SH3002803) and harvested by cell scraping. Cells were collected via centrifugation and lysed with lysis buffer (50 mM Tris, pH 7.5, 150 mM NaCl, and 2 mM n-Dodecyl-β-D-maltoside (DDM; GoldBio, catalog #: DDM5)), supplemented with complete protease inhibitor cocktail (Roche, catalog #: 4693159001). Cells were subjected to three rounds of vortexing and sonication for 1 min each, and the lysates were cleared by centrifugation (21,000×g, 10 min, 4 °C). Protein concentration was determined by MicroBCA assay kit (Thermo Fisher Scientific, catolog #: 23235). To prepare samples for SDS-PAGE, 4x Laemmli sample buffer (Bio-Rad, catalog #: 161-0747) with 10% β-mercaptoethanol (β-ME, Millipore Sigma catalog #: M3148) were added to protein samples. Protein samples were separated in an 8% SDS-PAGE gel. The gel was then transferred onto nitrocellulose membranes (Bio-Rad, catalog #: 162-0115), which were blocked with 5% milk in TBS-T (TBS with 0.1% Tween20 (Fisher Scientific, #BP337-500)) for 1 hour at room temperature. The blots were incubated with primary antibodies in 1% milk in TBS-T overnight at 4°C on shaker. On the second day, the blots were washed with TBS-T and incubated with secondary antibodies in 1% milk in TBS-T for 1 hour at room temperature. Finally, the blots were washed with TBS-T and developed using SuperSignal Pico PLUS or Femto chemiluminescent substrates (Fisher Scientific, catalog #: PI34578, catalog #: PI34095). Proteins were visualized using Azure Biosystems C600.

The following primary antibodies were used: the mouse monoclonal anti-GABA_A_α1 subunit antibody (1:2000; catalog #: MAB339), mouse monoclonal anti-GABA_A_β2/3 subunit antibody (1:1000, catalog #: 05–474), rabbit poly-clonal anti-GABA_A_β2 subunit antibody (1:1000, catalog #: AB5561), rabbit polyclonal anti-GABA_A_γ2 subunit antibody (1:1000, catalog #: AB5559), rabbit monoclonal anti-Na^+^/K^+^ ATPase antibody (1:10,000; catalog #: ab76020), mouse monoclonal anti-ubiquitin antibody (1:2000; catalog #: 14-6078-82), rabbit polyclonal anti-LC3b antibody (1:2000; catalog #: NB100-2220), and the fluorescent hFAB Rhodamine anti-β-actin antibody from Biorad (catalog #: 12004163). The following secondary antibodies were used: goat anti-Rabbit IgG Superclonal secondary antibody (1:10,000; catalog #: A27036) and goat anti-Mouse IgG Superclonal recombinant secondary antibody (1:10,000; catalog #: A28177), peroxidase AffiniPure goat anti-Mouse IgG secondary antibody, light chain specific (1:10,000, catalog #: 115-035-174), or peroxidase IgG monoclonal mouse anti-Rabbit IgG, light chain specific (1:10,000, catalog #:211-032-171).

### Cell surface protein biotinylation

HEK293T cells were seeded in poly-L-lysine (PLL)-coated 6-cm dishes according to previously published procedure (34) and subjected to transfection as described above. On the day of harvesting, intact cells were washed with ice-cold Dulbecco’s phosphate buffered saline (DPBS) (Fisher Scientific, catalog #: SH3002803). To label surface membrane proteins, cells were incubated with the membrane-impermeable biotinylation reagent Sulfo-NHS SS-Biotin (0.5 mg/mL; Pierce, catalog #: 21331) in DPBS containing 0.5 mM CaCl_2_ and 1 mM MgCl_2_ (DPBS-CM) for 20 min on ice. To quench the reaction, cells were incubated with 50 mM glycine in ice-cold DPBS-CM twice for 5 min on ice. Cells were then washed with DPBS twice. Sulfhydryl groups were blocked by incubating the cells with 5 nM N-ethylmaleimide (NEM; Pierce, catalog #: 23030) in DPBS for 15 min at room temperature. Cells were collected by scrapping and then solubilized in lysis buffer (50 mM Tris, pH 7.5, 150 mM NaCl, and 2 mM DDM), supplemented complete protease inhibitor cocktail and 5 mM NEM overnight on orbital rotator at 4°C. The following day, samples were centrifuged at 21,000×g for 10 min at 4 °C to pellet cellular debris. The supernatant contained the biotinylated surface proteins. Biotinylated surface proteins were affinity-purified from the above supernatant by incubating for 2 hr at 4 °C with 50 μL of immobilized neutravidin-conjugated agarose bead slurry (Pierce, catalog #: 29200). The beads were pelleted at 5000×g for 30 s and washed three times with wash buffer with detergent (TBS with 1% Triton X-100), and three times with wash buffer without detergent (TBS). Finally, surface proteins were eluted from beads by vortexing for 30 min with 80 μL of LSB ⁄ Urea buffer (2 x Laemmli sample buffer (LSB) with 100 mM DTT and 6 M urea; pH 6.8) for downstream SDS-PAGE and Western blotting analysis.

### Immunofluorescence and confocal microscopy

Immunofluorescence staining and confocal microscopy analysis were performed as described previously (34). Briefly, to label cell surface proteins, HEK293T cells transiently expressing WT or β2 variants on coverslips were fixed with 4% paraformaldehyde in DPBS for 15 min. For surface β2 detection, cells were not permeabilized. Fixed cells were blocked with 10% goat serum (Thermo Fisher, catalog #: 16210064) in DPBS for 30 min and incubated with 100 μL of mouse anti-GABA_A_ β2/3 subunit antibody (1:300, catalog #: 05–474) and rabbit anti-Na^+^/K^+^ ATPase antibody (1:300; catalog #: ab76020) in 2% goat serum in DPBS at 4°C overnight. For total protein staining, fixed cells were permeabilized with 0.2% saponin in DPBS and then blocked with 10% goat serum in DPBS for 30 min. Cells were incubated with 100 μL of mouse anti-GABA_A_ β2/3 subunit antibody (1:300, catalog #: 05–474) and rabbit anti-calnexin antibody (1:300; catalog #:ADI-SPA-860-F) in 2% goat serum and 0.2% saponin in DPBS at 4 °C overnight. The next day, cells were incubated in Alexa 488-conjugated goat anti-mouse antibody (Thermo Fisher, catalog #: A-11029) and Alexa 568-conjugated goat anti-rabbit antibody (Thermo Fisher, catalog #: A-11036) (1:500 dilution) in 2% goat serum with (total) or without (surface) 0.2% saponin in DPBS for 1 hr. Then, to stain the nucleus, cells were incubated with DAPI (1 μg/mL) (Thermo Fisher, catalog #: D1306) for 5 min. The coverslips were mounted using fluoromount-G (VWR, catalog #: 100502–406) and sealed with nail polish. An Olympus IX-81 Fluoview FV3000 confocal laser scanning system was used for imaging with a 60x 1.40 numerical aperture oil objective using the FV31S-SW software. The images were analyzed, and surface intensity was quantified using the ImageJ software.

### Automated patch clamping

Automated patch clamping electrophysiology was performed in HEK293T cells transiently expressing GABA_A_ receptors WT or β2 variants using the Ionflux Mercury 16 instrument (Fluxion Biosciences, California), as previously described(33). Briefly, cells grown to 90% confluency in 10-cm plates on the day of experiment were detached using accutase (Sigma Aldrich, catalog #: A6964) and suspended in serum free medium HEK293 SFM II (Gibco, catalog #: 11686–029) in the presence of 25 mM HEPES (Gibco, catalog #: 15630–080) and 1% penicillin streptomycin. Cells in suspension were placed on a shaker and left to shake for 30 min to 1 hr at room temperature. Then, cells were pelleted at 200×g for 1 min, resuspended in the extracellular solution (ECS; 138 mM NaCl, 4 mM KCl, 1 mM MgCl2, 1.8 mM CaCl2, 5.6 mM Glucose, 10 mM HEPES), and added to the Ionflux ensemble plate 16 (Fluxion Biosciences, catalog #: 910–0054). To determine peak chloride current, a saturating concentration of GABA (100 µM; Millipore Sigma, catalog #: A2129) was applied for 3 seconds. Whole-cell GABA-induced currents were recorded at a holding potential of –60 mV, and each ensemble recording enclosed the current of 20 cells. The signals were acquired and analyzed by Fluxion Data Analyzer.

### Cycloheximide (CHX) chase assay

Cycloheximide (100 μg/mL) was added to HEK293T cells transiently expressing GABA_A_ receptors WT or β2 variants at various time points (0-6 hrs) forty-eight hours post transfection to inhibit protein synthesis. Cells were harvested, lysed and ran on an 8% SDS-PAGE gel as previously described. Bands were normalized to loading control (β-actin) and further normalized to time point 0 to visualize the decay.

### Immunoprecipitation

Cell lysates (1000 μg) were pre-cleared with 30 μL of protein A/G plus-agarose beads (Santa Cruz Biotechnology, catalog #: SC-2003) and 1 μg of normal mouse IgG (Santa Cruz Biotechnology, catalog #: SC-2025) or normal rabbit IgG (Santa Cruz Biotechnology, catalog #: SC-2027) for 1 hr at 4 °C to remove nonspecific binding proteins. The pre-cleared samples were incubated with 2 μg of mouse anti-GABA_A_α1 antibody, mouse anti-GABA_A_β2/3 antibody, rabbit anti-GABA_A_β2 antibody, rabbit polyclonal anti-GABA_A_γ2 subunit antibody or normal mouse/rabbit IgG as a negative control for 1 hr at 4 °C. The protein-antibody complexes were then incubated with 50 μL of protein A/G plus agarose beads overnight at 4 °C. The next day, the beads were collected by centrifugation at 3000×g for 30 s. Supernatant, which contained non-interacting proteins, was discarded. Beads were washed three times with wash buffer (TBS with 2 mM DDM) and eluted by incubation with 60 μL of 2x Laemmli sample buffer with 10% β-ME, boiling at 42°C for three min and vortexing for 10 min. The immunopurified eluents were separated in an 8% SDS-PAGE gel, followed by western blot analysis using appropriate antibodies.

### Endoglycosidase H (Endo H) assay

HEK293T cells transiently expressing GABA_A_ receptors WT or β2 variants were harvested and lysed as described previously. To denature the protein, cell lysate (30 µg) was incubated with Glycoprotein Denaturing Buffer for 10 min at room temperature. Then, endoglycosidase H enzyme (endo H; NEBiolab, catalog #: P0703L) and GlycoBuffer were added to the denatured solution for 2 hrs at 37°C to digest the glycoproteins. Undigested samples where no endo H was added were used the negative control. Peptide-*N*-Glycosidase F (PNGase F, NEBiolab, catalog #: P0704L), which cleaves almost all glycans, were added to the denaturation solution with GlycoBuffer and NP40 as a positive control. After the 2-hr incubation, 4x Laemmli sample buffer with 10% β-mercaptoethanol was added to the samples, which were ran on an 8% SDS-PAGE gel for western blot analysis.

### Quantitative reverse transcription polymerase chain reaction (qRT-PCR)

HEK293T cells transiently expressing GABA_A_ receptors WT or β2 variants were harvested forty-eight hours post-transfection as described previously. Total RNA was extracted with the Roche High Pure RNA Isolation Kit (catalog #: 11828665001) and 1 µg of total RNA was converted to cDNA using the iScript^TM^ cDNA synthesis kit (Bio-Rad, catalog #: 1708891). For the polymerase chain reaction, cDNA was mixed with PowerUp SYBR Green Master Mix (Applied Biosystems, catalog #: A25776) and appropriate primers and amplified (40 cycles of 15 s at 95 °C and 60 s at 60 °C) using the QuantStudio 3 Real-Time PCR System (Applied Biosystems). The forward and reverse primers for genes of interest and GAPDH (housekeeping gene control) are listed in **Supplementary Table S1**.

Amplification results were analyzed with the QuantStudio software, and the threshold cycle (C_T_) was obtained from the PCR amplification plot. ΔC_T_ value was calculated as:

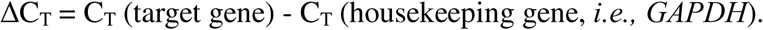

The relative mRNA expression level of target genes of variant was normalized to WT cells:

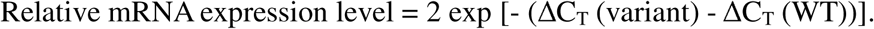

Each data point represents average from three technical triplicates.

### Statistical analysis

Quantification of band and pixel intensity was performed using ImageJ. Data was plotted and statistical significance was determined using GraphPad Prism. All data are presented as mean ± SEM. If two groups were compared, statistical significance was calculated using an unpaired Student’s t-test; if more than two groups were compared, ANOVA followed by post hoc Dunnett’s test was applied. A *p*<0.05 was considered statistically significant. *, *p*<0.05; **, *p*<0.01; ***, *p*<0.001; ****, *p*<0.0001.

## Supporting information

This article contains supporting information. Supporting information includes one supplemental table and two supplemental figures.

## Acknowledgements

This work was supported by the National Institutes of Health (R01NS105789 and R01NS117176 to TM) and American Heart Association pre-doctoral fellowship (25PRE1372186 to XC).

## Author contributions

Conceptualization, XC, YW, and TM; Data curation: XC; Formal analysis: XC, YW, and TM; Funding acquisition: XC and TM; Supervision: YW and TM; Writing – original draft: XC, YW, and TM; Writing – review & editing: XC, YW, and TM.

## Supplemental Table

**Table S1.**
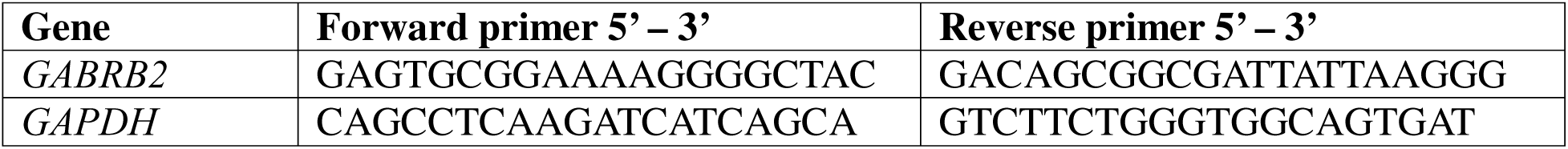
List of qPCR primers that were used in this work.

## Supplemental figure legends

**Supplemental Figure S1.**
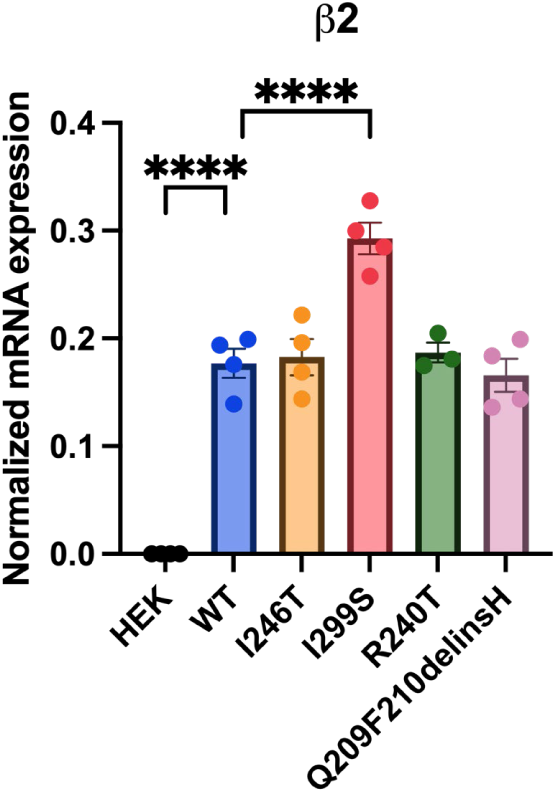
mRNA levels of WT and. β**2 EAVs.** HEK293T cells transiently expressing WT or β2 EAVs were harvested forty-eight hours post transfection. Total RNA was isolated and converted to cDNA for quantitative PCR amplifications to detect the relative β2 expression. HEK293T cell without transfection was included as a negative control. GAPDH was used as a house-keeping gene. Each gene was analyzed in triplicates and the experiment was performed three times (n=3). Data is presented as mean ± SEM. One-way ANOVA followed by a post-hoc Dunnett’s test was used to determine statistical significance. ****, *p* < 0.0001.

**Supplemental Figure S2.**
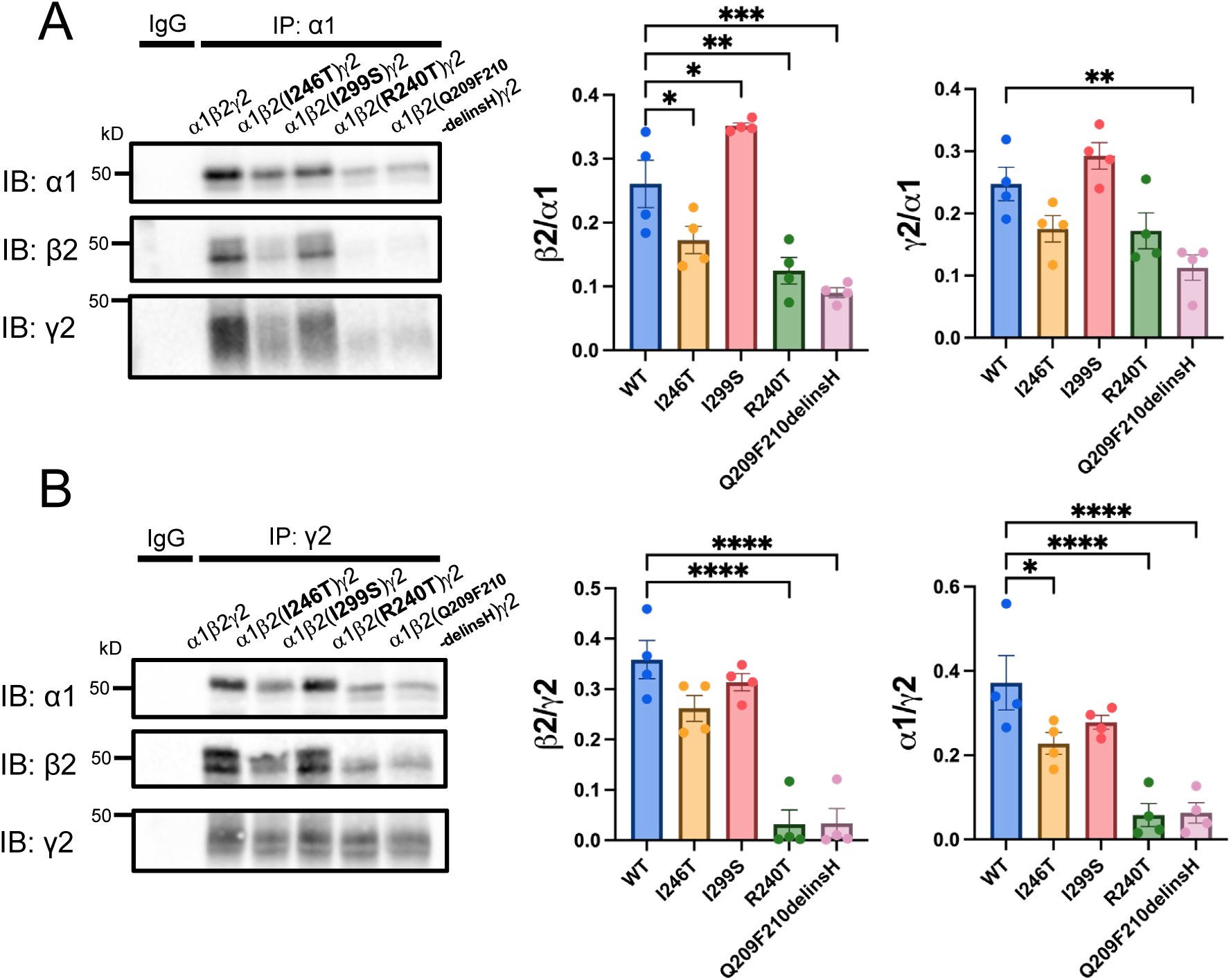
Interaction between. α**1,** β**2, and** γ**2 subunit of** β**2 EAVs.** (**A**) Interaction between β2 and α1, and between γ2 and α1 subunits. Co-IP experiment was performed with α1 as the bait protein. Quantification of the ratio of proteins of interest (i.e., β2 or γ2) to α1 is shown in the right panels (n=4). (**B**) Interaction between β2 and γ2, and between α1 and γ2 subunits. Co-IP experiment was performed with γ2 as the bait protein. Quantification of the ratio of proteins of interest (i.e., β2 or α1) to γ2 is shown in the right panels (n=4). IP: immunoprecipitation. Data is presented as mean ± SEM. One-way ANOVA followed by a post-hoc Dunnett’s test was used to determine statistical significance. * *p*< 0.05; **, *p* < 0.01; ***, *p* < 0.001; ****, *p* < 0.0001.

